# CryoEM structure of ALK2:BMP6 reveals distinct mechanisms that allow ALK2 to interact with both BMP and Activin ligands

**DOI:** 10.1101/2025.02.07.637115

**Authors:** Erich J. Goebel, Senem Aykul, Kei Saotome, Aris N. Economides, Matthew C. Franklin, Vincent J. Idone

**Affiliations:** Connective Tissue Diseases Therapeutic Focus Area, Regeneron Pharmaceuticals, Tarrytown, NY 10591; Structural Biology, Regeneron Pharmaceuticals, Tarrytown, NY 10591

**Author notes:** **Corresponding Author**: Erich J. Goebel, 66 Woodbine Drive, Mahopac, NY, 10541. **Classification**. Biological Sciences/Biochemistry.

**Keywords:** TGF-Beta Family, Structural Biology, Skeletal Biology, Bone Morphogenetic Protein, Activin

## Abstract

Ligands in the TGFβ family (activins, BMPs and TGFβs), signal by bringing together two type I and two type II receptors. ALK2 is the only type I receptor among the seven TGFβ type I receptors that interacts with both activin and BMP ligands. With BMPs, ALK2 acts as a signaling receptor to activate SMAD1/5/8 signaling. Alternatively, with activins, such as Activin A (ActA), ALK2 forms nonsignaling complexes that negatively regulate ALK2 and ActA signaling. To gain insight into how ALK2 interacts with two distinct classes of ligands, we resolved the cryoelectron microscopy structure of ALK2 in complex with the type II receptor, ActRIIB, and the ligand, BMP6, in parallel with the corresponding structure with ALK3 for direct comparison. These structures demonstrate that ALK2 and ALK3 utilize different mechanisms to interact with BMP6 at the wrist interface, with ALK2 relying on BMP6 glycosylation and ALK3 relying on a salt bridge. Modeling of ALK2:ActA reveals that binding relies on ActA’s fingertip region, mirroring the interaction of ActA with its other receptor, ALK4. Our results demonstrate that ALK2 is a ‘hybrid’ receptor that incorporates features of BMP type I receptors such as ALK3 at the wrist interface, and an activin type I receptor such as ALK4 at the fingertip.

**Significance:** ALK2 interacts with two different classes of TGFβ ligands: the BMPs and the activins. With BMP ligands, ALK2 signals to play roles in bone modeling, while with activins, ALK2 negatively regulates signaling by forming nonsignaling complexes. We demonstrate that ALK2 binds to favored ligand, BMP6, by utilizing the ligand wrist-helix stabilized by a glycosylation, rather than a charged interaction observed with similar receptor, ALK3. We also show that ALK2 interaction with activin ligand, ActA, is reliant on a single interaction at the other side of the receptor – the fingertip interface. Thus, this study elucidates first structure of ALK2 in complex with a ligand and provides the molecular insight into how ALK2 is able to bind to BMP and activin ligands.

## Introduction

The transforming growth factor β (TGFβ) family is comprised of over 30 structurally similar ligands that fall into three major classes, each defined by downstream SMAD activation profiles and receptor specificity (1). Signaling is initiated by the formation of complexes comprised of one dimeric ligand, two type II receptors and two type I receptors with each ligand class using distinct mechanisms of complex formation(2). With five type II and seven type I receptors in mammals, there is a receptor bottleneck where an elaborate web of specificity, affinity, and competition has evolved to govern ligand:receptor interactions (1, 3). Bone Morphogenetic Proteins (BMPs), which contains more than 20 members, is the largest of the classes and activates the SMAD1/5/8 pathway through three type II receptors (ActRIIA, ActRIIB, BMPR2) and four type I receptors (ALK1/ACVRL1, ALK2/ACVR1, ALK3/BMPR1a and ALK6/BMPR1b). Each ligand:receptor combination possesses distinct signaling effects resulting from the cell-specific receptor repertoire, ligand availability as well as binding potency (3). ALK2 and its ligands present an example of this observation in that the response of ALK2 to BMPs can be negatively modulated by Activin A (ActA) both *in vitro* and *in vivo* (4, 5).

Interestingly, initial studies identified ALK2 (Activin receptor type 1) as an activin type I receptor (6, 7). However, further experiments revealed that ActA does not activate ALK2 whereas BMP ligands, such as BMP6 and BMP7, do (8). Hence, ALK2 was reclassified as a BMP receptor and the observed interaction with ActA was considered artifactual. This view was later revised when it was shown that in the genetic disorder, fibrodysplasia ossificans progressiva (FOP), single amino acid changes in the intracellular domain of ALK2 render the receptor responsive to ActA (4, 9). Subsequently, we showed that wildtype ALK2 forms non-signaling complexes with ActA and associated type II receptors (4, 5, 10) and that these non-signaling complexes traffic to the lysosome wherein their components are degraded (11). These findings reestablished ALK2 as an Activin receptor and indicated that ALK2 has evolved the unique capability to interact with two distinct class of ligands: the Activins and the BMPs.

To date, there is no published structure of ALK2 in complex with a ligand. This is likely due to difficulties not only producing active recombinant receptor for structural studies but validating the receptor’s proper folding and activity as well. Previous efforts to determine binding kinetics for ALK2 have yielded only mild success for several BMP ligands(5). This is in direct contrast to the nanomolar affinity seen between the other BMP type I receptors such as ALK1 and ALK3(12–14). Furthermore, the ligand with the highest affinity for ALK2, BMP6, is reliant on N-linked glycosylation to bind(15). Interestingly, whereas this post-translational modification on BMP6 is necessary for interaction with ALK2, it is dispensable for ALK3 or ALK6, suggesting differences in the manner of type I engagement.

To explore, at the molecular level, how ALK2 interacts with its cognate ligands, we utilized cryoelectron microscopy (cryoEM) to resolve the ternary complex structures of ALK2:ActRIIB:BMP6 and ALK3:ActRIIB:BMP6. Comparisons of these structures, along with biochemical validation, illustrate two mechanisms for stabilizing type I:BMP6 interactions which dictate the ability to bind to both ALK2 and ALK3. Extending our analysis of ALK2 through modeling and mutagenesis, we pinpoint a key residue in ALK2 that is critical for interaction with ActA, but not BMP6.

## Results

### ALK2-ActRIIB and ALK3-ActRIIB bind BMP6 with high affinity

To alleviate potential concerns with the low ligand affinity of ALK2, we designed fusion molecules featuring ALK2 tethered to the type II receptor, ActRIIB, as the latter generally has higher affinity for BMP ligands than ALK2 (5). This strategy of tethering the lower affinity type I receptor to the higher affinity type II receptor has been used previously to capture low-affinity interactions as well as for engineering ligand-blocking therapeutics, commonly referred to as ligand traps (16–19). For this study, the extracellular domains (ECDs) of human ALK2 (1-123) or ALK3 (1-152) and ActRIIB (1-133) were fused to IgG1 Fc domains (SFig. 1). ALK2-ActRIIB-Fc and ALK3-ActRIIB-Fc were generated through co-expression and purified to homogeneity prior to biochemical validation (SFig. 1).

**Figure 1.**
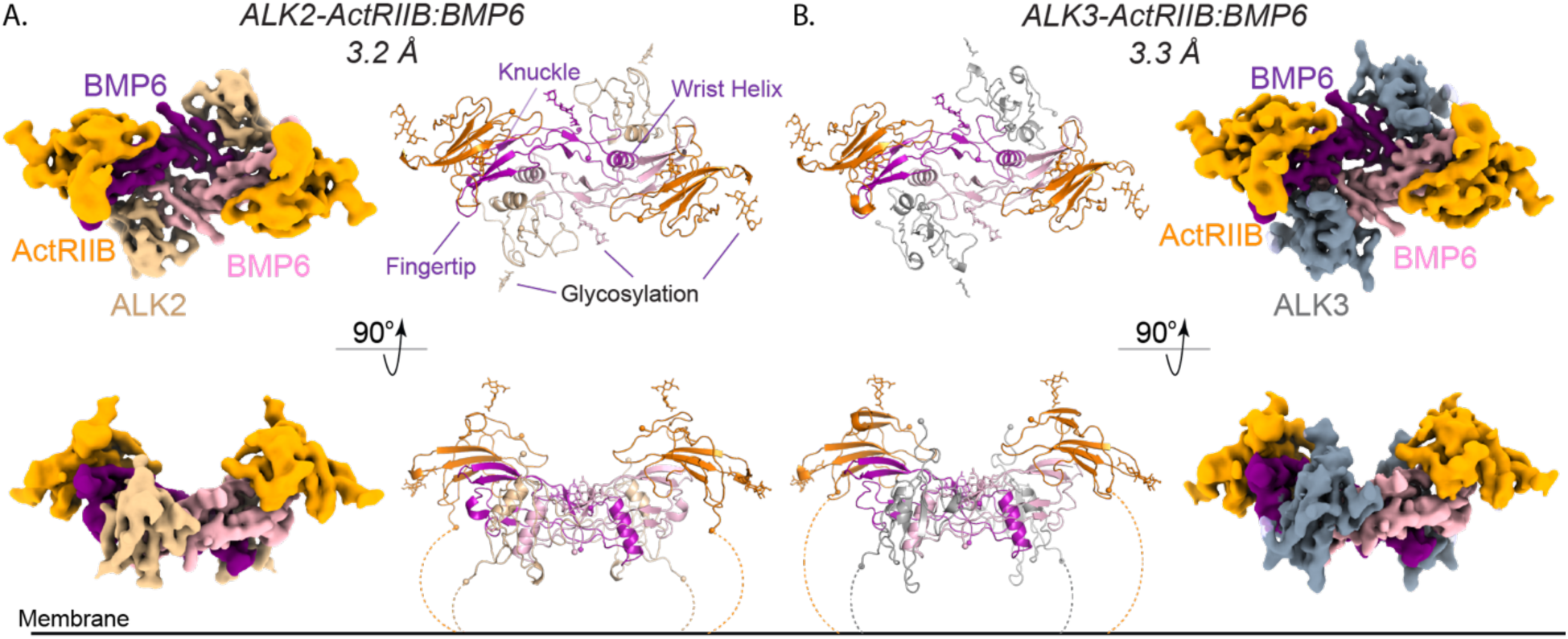
CryoEM structures of BMP6 in complex with ActRIIB-ALK2 and ActRIIB-ALK3. **(A. and B.)** Two views of the 3.23 Å and 3.3 Å resolution cryoEM maps and resulting structures of BMP6 in complex with ActRIIB-ALK2 **(A.)** and ActRIIB-ALK3 (**B.)**, respectively. ALK2 (*wheat*) and ALK3 (*gray*) bind in the composite Type I interface built from both BMP6 monomers (*purple/pink*). ActRIIB (*orange*) binds at the ligand knuckle interface, dependent on a single monomer.

To determine if the receptor ECDs within our traps were properly folded, we utilized surface plasmon resonance (SPR) to kinetically measure the binding of both heterodimeric traps to mammalian expressed (glycosylated) BMP6 (SFig 2). Additionally, we measured binding of two additional traps: one with a single ActRIIB ECD and another containing two ActRIIB ECDs (ActRIIB_1_-Fc and ActRIIB_2_-Fc respectively). Comparison of the affinities of the different traps revealed that the heterodimeric traps displayed a higher affinity for BMP6 (36.0pM; ALK2:ActRIIB and 51.7pM; ALK3:ActRIIB) than either ActRIIB_2_-Fc (123.6pM) or ActRIIB_1_-Fc (281.7pM) (SFig. 2 and STable 1). In corroboration with the SPR results, we were able to form stable complexes between both traps and BMP6 as determined by size exclusion chromatography (SEC) (SFig. 1). These data indicated that, in both heterodimeric traps, each of the receptor ECDs is active and binding to BMP6.

**Figure 2.**
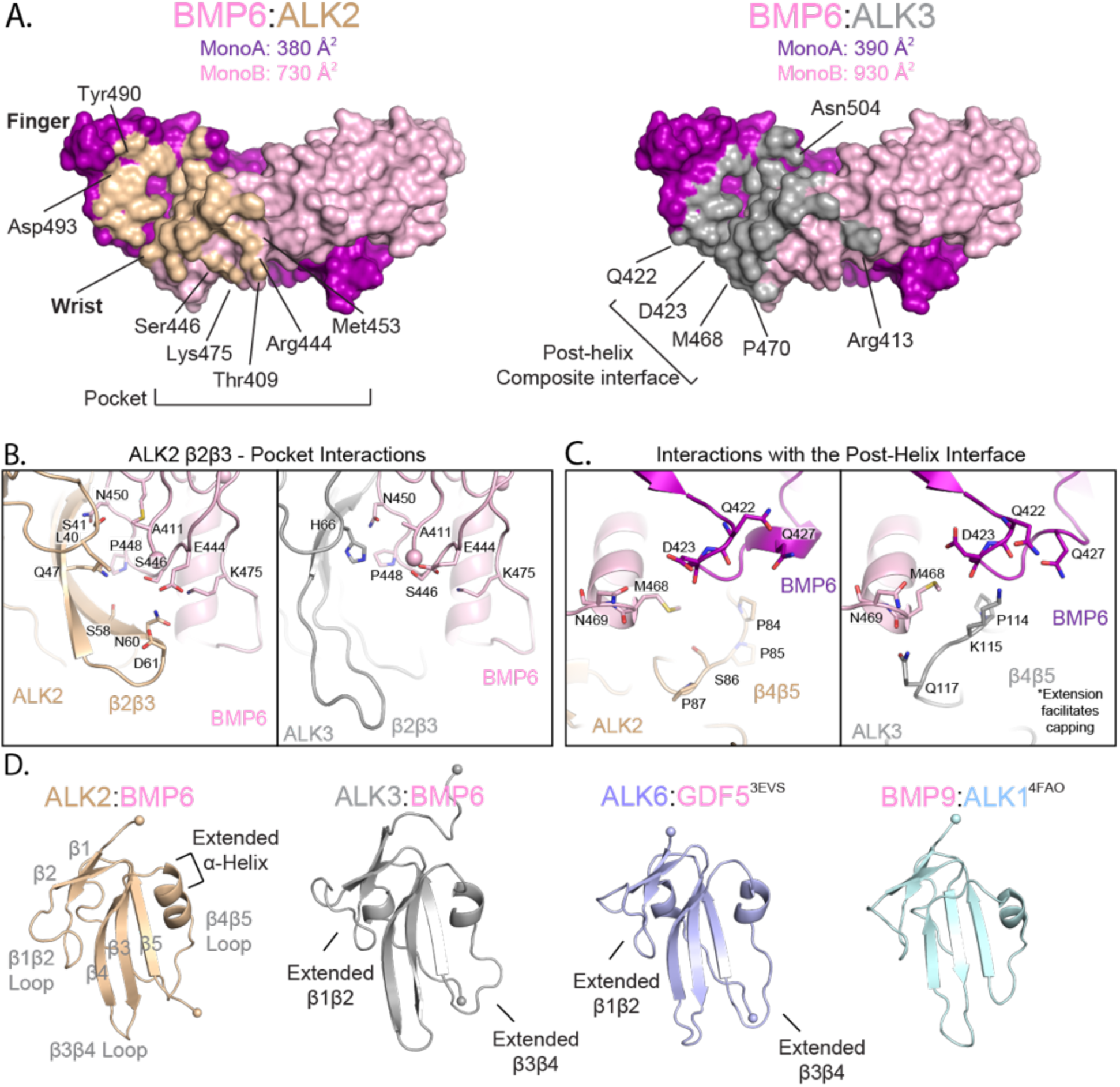
ALK2 and ALK3 utilize distinct interfaces to bind BMP6. **A.** Ligand surface representation of the Type I interface, with BMP6 residues interacting with either ALK2 or ALK3 colored in *wheat* and *gray* respectively. Buried surface area calculations for each of the interfaces are shown following determination in PISA. **B.** Comparison of the β2β3 loop interface during interaction with ALK2 (*left*) and ALK3 (*right*). **C.** Comparison of interactions between the post-helical region of BMP6 with ALK2 (*left*) and ALK3 (*right*). **D.** Comparison of ligand bound BMP type I receptors: BMP6 bound ALK2 and ALK3, GDF5 bound ALK6 (*blue*, PDB: 3EVS (27)) and BMP9 bound ALK1 (*light blue:* 4FAO (14)).

### CryoEM Structures of ALK2-ActRIIB and ALK3-ActRIIB bound to BMP6

We next sought to resolve the structure of ALK2-ActRIIB-Fc:BMP6. Initial cryoEM data collection of the complex revealed that the Fc portion of the trap was interfering with the resolution of the structure. Removing the Fc portion of the heterodimeric traps through IDES digestion had no significant impact on the ability of ALK2-ActRIIB and ALK3-ActRIIB, to form stable complexes with BMP6 (SFig. 1). Data collection on this smaller, ALK2-ActRIIB:BMP6 complex, yielded a high-resolution structure containing a complete complex with two ActRIIB-ECD and two ALK2-ECD bound. The cryoEM structure of ALK2-ActRIIB:BMP6 was resolved to 3.2Å and the map was well defined for residues: ALK2 (31-109), ActRIIB (26-118) and the mature BMP6 ligand (410-513; 82.3% of mature domain) (Fig. 1, Sfig. 3, STable 2). We used a similar experimental setup to resolve the structure of ALK3-ActRIIB bound to BMP6 to 3.3Å with residues, ALK3 (50-141), ActRIIB (26-117) and BMP6 mature domain (410-513), being well-resolved. In both structures, BMP6 adopts the canonical dimeric fold of TGFβ ligands, akin to ‘two hands’: two monomeric molecules held together by a cystine knot, where each contains two long β-ribbons of two antiparallel β-strands each (fingers) and a large alpha-helix (wrist) (Fig. 1)(1, 20). The overall complex assembly of ligand:receptors is consistent with previous BMP complex structures, with the type II receptor binding at the convex, ‘knuckle’ of the β-ribbons and the type I receptor binding at the interface built from the concave fold of the fingers and the wrist helix(21).

**Figure 3.**
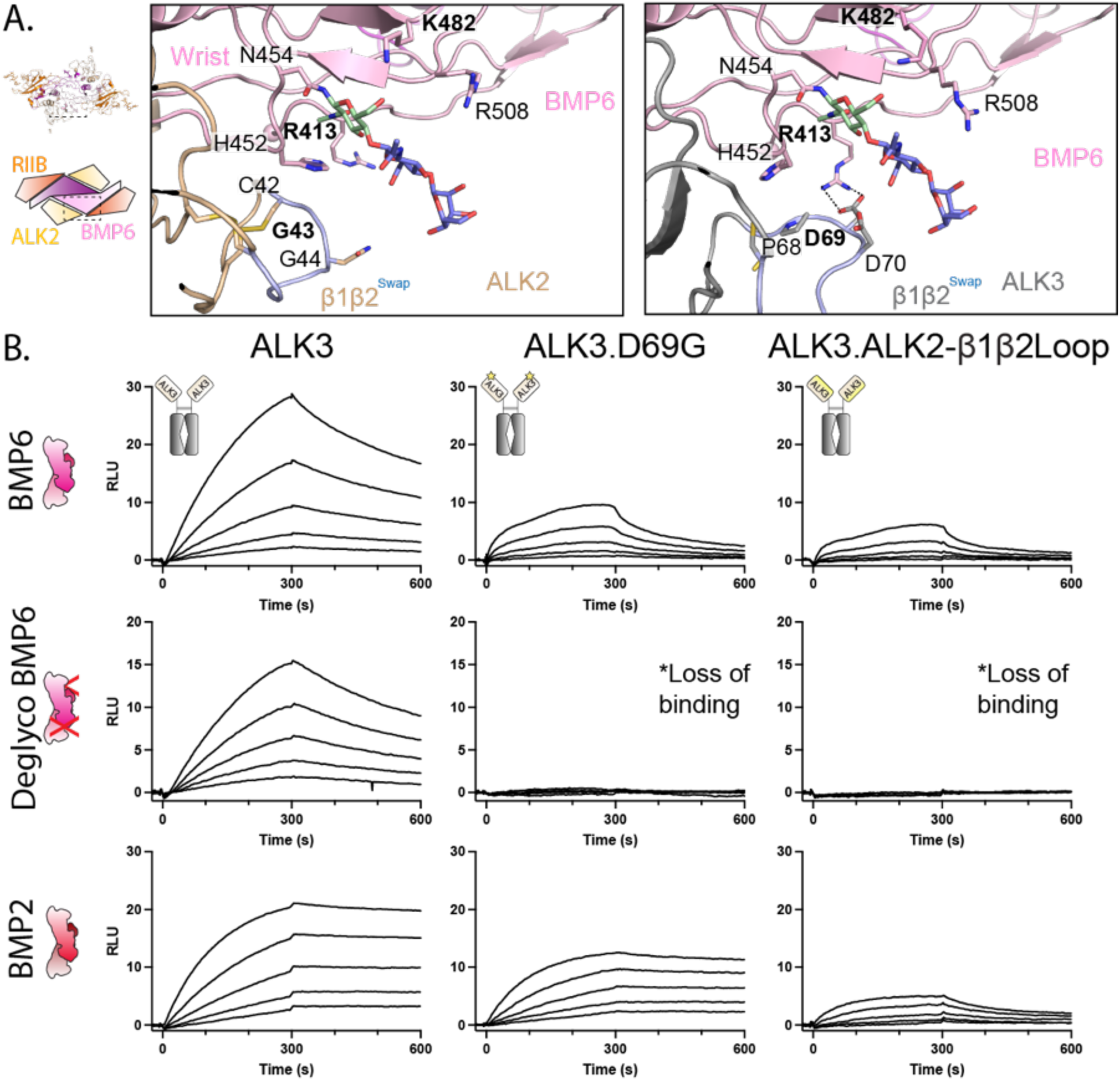
BMP6 utilizes two compensatory mechanisms to stabilize type I binding. **A.** Schematic view of ALK2-ActRIIB:BMP6 with comparison of the β1β2 interface, highlighting the position of the N-linked glycosylation (primary NAG in *green*; Second NAG and BMA in *blue*) on the BMP6 pre-helix and the formation of a salt bridge at the same interface between ALK3:BMP6. Engineered β1β2 loop swap between ALK2 and ALK3 colored in *light blue* in both structures. **B.** SPR sensorgrams of BMP6, deglycosylated BMP6, and BMP2 binding to anti-human antibody-captured ALK3-Fc, ALK3.D69G-Fc, and ALK3.ALK2-β1β2Loop-Fc. Each experiment was performed in duplicate.

Similar to previous structural efforts, we were unable to confidently build the antibody linker between the type II and type I arms of either ALK2-ActRIIB or ALK3-ActRIIB, likely due to flexibility (19). However, when contoured to a low level, the cryoEM maps suggest that the antibody linker is between the cis-pair of receptors and that the two heteromeric traps bind to the ligand in a cis-manner (SFig. 4). Measurements between the terminal tails for the different receptor pairs is consistent with what has been previously reported (∼35Å for the cis-pair and ∼70Å for the trans-pair) (22).

**Figure 4.**
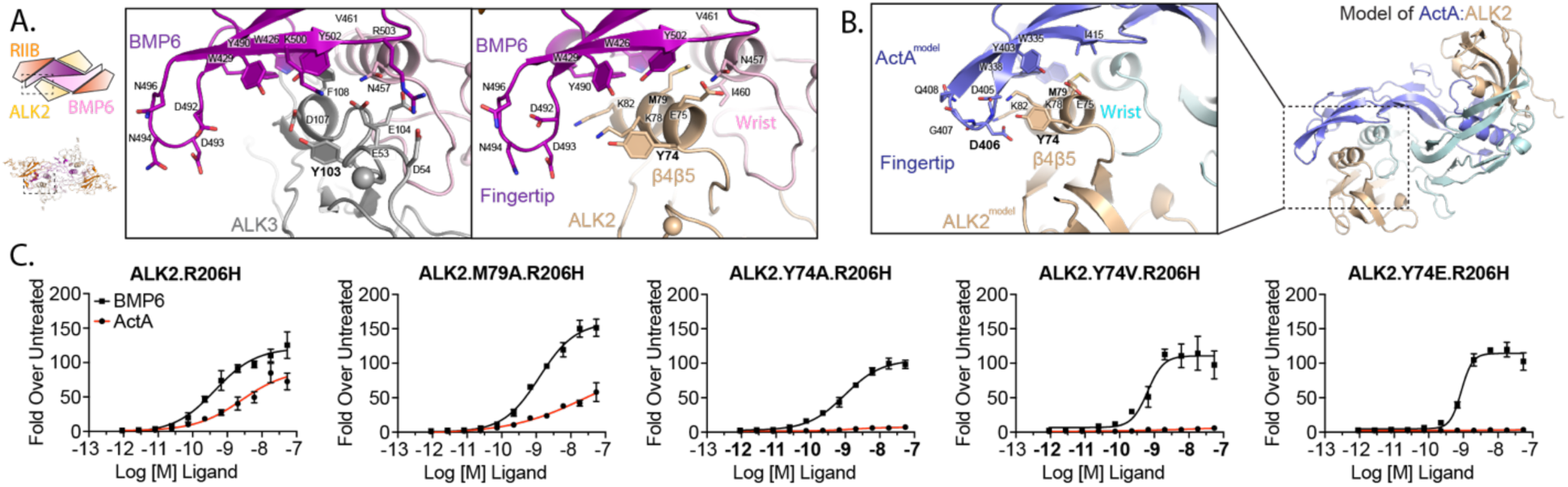
ALK2 uniquely engages the ActA fingertip for interaction and signaling. **A.** Schematic view of ALK2-ActRIIB:BMP6 with comparison of the fingertip interface between BMP6 bound ALK2, ALK3 **B.** AlphaFold model of the ALK2:ActA complex highlighting fingertip interface. **C.** Luciferase reporter assays in expression HEK cells, stably expressing either ALK2.R206H or respective variants. Cells were treated with a titration of BMP6 (*black lines*) and ActA (*red lines*).

### BMP6 binding to ActRIIB, ALK2 and ALK3

Both structures indicate that ActRIIB binds at the knuckle of BMP6, utilizing a highly conserved set of interfacial residues observed across numerous type II complex structures containing both BMP and Activin class ligands(20). Specifically, a hydrophobic triad (Tyr^60^, Trp^78^, and Phe^101^) in ActRIIB engages an “IAP” motif on BMP6 (Ile^22^, Ala^23^, and Pro^24^) to form the core of the receptor interface. The exact receptor pair (ALK2-ActRIIB or ALK3-ActRIIB) of the trap has little effect on ActRIIB during BMP6 binding (RMSD = 0.632 over 92 Cα atoms), indicating there are no significant constraints induced by the tethered type I receptor.

In both structures, the type I receptor binds within the concave, composite interface built from both ligand monomers, albeit with different surface area utilization. During interaction with ALK3, BMP6 buries a total of ∼1325.4 Å^2^, similar to that of ALK3:BMP2 (23), while only burying 1,113.8 Å^2^ while engaging ALK2. Comparison between the two structures not only reveals that ALK2 engages a much smaller degree of surface area (203.6 Å^2^ less) on BMP6 monomer B, but that the pattern of surface engagement is significantly different with each type I receptor engaging unique interfaces (Fig. 2). Notably, the β2β3 loop of ALK2 contacts a charged pocket on BMP6 forming several charged/polar interactions between several ALK2 (Ser^41^, Asn^60^, Asp^60^) and BMP6 (Asn^450^, Ser^446^, Glu^444^, Lys^475^) residues (Fig. 2B). The ALK3 β2β3 loop is positioned distantly and in fact, an overlay of the two structures shows a 27° shift of overall receptor positioning in the concave ligand cleft. Conversely, ALK3 caps the wrist helix with residue Gln^117^, engaging Asn^469^ of the post-helical loop of BMP6, whereas ALK2 does not (Fig. 2C). This interaction is facilitated by a three amino acid extension in the β4β5 loop of ALK3, not present in ALK2 (Fig. 2D). Further comparison of ALK2 and ALK3 reveals that the alpha helix that is formed in the ALK2 β4β5 loop is uniquely extended an additional half turn when compared to that of ALK3, as well as other BMP type I receptors, ALK6 or ALK1 (Fig. 2D). Despite this conformational difference, no major contacts are observed between this alpha helix extension and BMP6.

### Glycosylation of the BMP6 prehelical loop stabilizes interaction with ALK2

The differences in the manner of engagement of BMP6 by ALK2 versus ALK3 is accentuated further by the fact that only glycosylated BMP6 can activate ALK2 (15, 24). *In vivo*, BMP6 exists as several species corresponding to glycosylated and nonglycosylated forms (25, 26). Glycosylation of the prehelix loop determines whether BMP6 can activate ALK2; enzymatically deglycosylated or bacterially produced BMP6 (which is not glycosylated) is unable to bind within SPR experiments or signal through ALK2 (15, 24). We corroborate these previous findings by showing that deglycosylated BMP6 (dgBMP6) does not interact with ALK2 on SPR (SFig. 5). Here, we show that BMP6 binds ALK2-ActRIIB-Fc with a KD of 36pM whereas dgBMP6 has a reduced KD of 164.3pM (SFig 2 and 5, STable 1). Conversely, deglycosylation of BMP6 does not significantly reduce binding to ALK3-ActRIIB-Fc to the same degree (KD of 51.7pM and 71.4pM for BMP6 and dgBMP6, respectively). Additionally, several structures of BMP ligands, particularly BMP2, show that the prehelix loop is flexible but is stabilized in a conserved conformation upon type I binding (14, 23, 27–31). Given the requirement for glycosylation along with the flexibility of the loop that contains it, we utilized the structure of ALK2-ActRIIB:BMP6 to explore the carbohydrate’s role in ALK2 binding.

The cryoEM structure of ALK2-ActRIIB:BMP6 displays a strong signal for the first three carbohydrate moieties of the prehelical Asn^454^–linked glycosylation (2x N-acetylglucoasmine and 1x β-d-mannose). Surprisingly, no molecular contacts are seen between carbohydrate chain of BMP6 and ALK2, suggesting that there is no direct contribution of the carbohydrate chain to ALK2 binding (Fig. 3A). Notably, previously generated structural models of ALK2 bound to BMP6 suggest that residues Lys^31^ and Tyr^74^ directly interact with the glycosylation (15), whereas our structure reveals that both residues are positioned on the opposite side of the receptor and away from the wrist interface, hence precluding direct binding (SFig. 6) (15). We do observe, however, several interactions between the carbohydrate chain and specific residues in the same monomer chain of BMP6. Specifically, the primary carbohydrate moiety forms a stabilizing hydrophobic interaction with His^452^, as well as forming a hydrogen bond with Lys^482^ (Fig. 3A). These two interactions stabilize BMP6’s prehelix loop into a conformation suitable for ALK2 binding and mirroring previous structures of type I bound BMP ligands. Along this idea, no significant structural differences are observed in the backbone chain of BMP6 when comparing the ALK2 and ALK3-bound structures (RMSD = 0.624 Å over 208 Cα atoms), supporting the overall rigidity of BMP6 and the necessity for a specific prehelix loop conformation.

Given the lack of direct binding between the carbohydrate moiety of BMP6 and ALK2, it remains curious why ALK2 is reliant on its presence for BMP6 binding, whereas ALK3 is not. Comparison between the CryoEM structure of ALK2:BMP6 and ALK3:BMP6 reveals the presence of a salt bridge between ALK3 residue Asp^69^ and BMP6 residue Arg^413^, which sterically occludes the previously mentioned His^452^ driven hydrophobic contacts in BMP6 (Fig. 3A). The β1β2 loop in ALK2 is three residues shorter and a glycine residue is in the position of ALK3 Asp^69^ preventing a similar contact. Additionally, Arg^413^ is positioned away from ALK2, further suggesting no interaction. We hypothesized that ALK3 utilizes this salt bridge to stabilize the prehelix loop without relying on the carbohydrate moiety.

To investigate this further, we produced and purified the homodimeric Fc-fusion of ALK3-ECD, and a corresponding version where Asp^69^ is replaced with the homologous ALK2 residue, a glycine (ALK3.D69G). We also generated an ALK3 ECD variant where the entire β1β2 loop (from cysteine-cysteine to preserve disulfide bonds) was replaced with that of ALK2 (ALK3.ALK2-β1β2Loop) (Fig. 3A). The effects on ligand specificity and binding were determined through SPR binding studies with the receptor fusions captured on the biosensor via the Fc domain (Fig. 3B). Binding was observed between ALK3-Fc and both BMP6 and dgBMP6. However, significantly reduced binding of dgBMP6 was observed with ALK3.D69G. Similar effects were seen during binding to ALK3.ALK2-β1β2Loop. In contrast, BMP2 binding was similar between ALK3 and ALK3.D69G, where only introduction of the β1β2 from ALK2 reduces binding significantly (Fig. 3B). This data supports that the salt bridge between ALK3 Asp^69^ and BMP6 Arg^413^ stabilizes binding and constitutes an alternative strategy to stabilize the prehelix loop in the absence of glycosylation. In contrast, ALK2, with a glycine in this position is unable to form the salt bridge with BMP6 and is thus reliant on the presence of the glycosylation to stabilize the pre-helical loop’s conformation for binding.

### The binding between ActA and ALK2 is reliant on unique fingertip:β4β5 interactions

Activin ligands, such as ActA or GDF11, uniquely rely heavily on their fingertips for type I receptor affinity as well as specificity, where extensive hydrogen bond networks are formed between residues within the ligand fingertip and the β4β5 of the type I receptor (19, 22). While a structure of ActA in complex with ALK2 proved difficult to resolve, we can draw insight from the structure of ALK2 bound to BMP6 for possible mechanisms of ALK2:ActA binding.

Analysis of the fingertip:β4β5 interface in our structures reveals that no direct interactions occur between the fingertip of BMP6 and ALK2 (Fig. 4A). This is consistent with previously reported type I:BMP structures, including our cryoEM structure of ALK3:BMP6. However, the additional helical turn in the β4β5 loop of ALK2 positions the C alpha carbon of Tyr^74^ 3.8 Å closer to the ligand fingertip than the corresponding Tyr^103^ residue in ALK3 (Fig 4A). Next, we aligned both the structure of ALK4:ActA (PDB: 7OLY) to ALK2:BMP6 (SFig. 7) as well as utilized AlphaFold to model ALK2:ActA (Fig. 4B) in order to visualize whether this shift in ALK2’s relative β4β5 position could facilitate unique contacts with ActA (32). In both strategies we observed that Asp^406^ of the ActA fingertip is well-positioned to form a hydrogen bond with Tyr^74^ of ALK2. Thus, we hypothesized that ALK2 is able to bind to ActA due to the unique positioning and conformation of the β4β5 loop and specifically residue Tyr^74^.

To probe this specific contact and its possible role on ActA:ALK2 interaction, we generated a set of stable cell lines expressing various ALK2 mutants starting with parental cell line expressing a BMP signaling reporter: three variations of Tyr^74^ as well as a mutation of knob-in-hole residue, Met^79^. Given that wildtype ALK2 is not activated by ActA, we utilized the FOP variant, ALK2.R206H in our construct design and thus, were able to utilize Smad1/5/8 signaling as a readout. As expected, both ActA and BMP6 activated ALK2.R206H cells (Fig. 4C). When Tyr^74^ was mutated to neutral residues, alanine or valine, ActA signaling was effectively ablated, while having minimal effect on BMP6 signaling via ALK2 (Fig. 4C). Substituting Tyr^74^ of ALK2 to the corresponding residue in ALK3, glutamate, ALK2 also lost the ability respond to ActA, but not BMP6. This corroborates with our previous work that modulated Asp^406^ in ActA and observed significant reduction in the ability of ActA to dimerize ALK2 and type II receptor (5). Unexpectedly, altering the knob-in-hole residue, Met^79^, to alanine did not ablate the signaling of either ligand, indicating that ALK2 relies more heavily on interactions outside of the conserved anchor interface to stabilize ligand binding. These results support our structural modeling and indicate that ActA engages ALK2 specifically with the ligand fingertip, relying on the unique structure of the ALK2 β4β5 loop and residue, Tyr^74^.

## Discussion

ALK2 is a type I receptor that binds two different ligand classes of the BMP/TGFβ superfamily: the BMPs and the Activins. To investigate how ALK2 accomplishes this task, we solved the cryoEM structures of both ALK2-ActRIIB and ALK3-ActRIIB bound to BMP6. Comparison of these structures reveals that although the BMP6 prehelix adopts the same conformation in both structures, the mechanism employed for stabilizing this shape of the prehelix is distinct between ALK2 and ALK3. With ALK2, BMP6 utilizes a glycosylation on Asn 454 to form interchain contacts that stabilize the prehelix in the active conformation, whereas with ALK3, BMP6 utilizes a salt bridge to stabilize the same interface.

The requirement for BMP6 to be glycosylated to activate ALK2 had been previously recognized (15). Our results illuminate the molecular details of this requirement but also hint to a potential regulatory role of BMP6 glycosylation. Glycosylated and deglycosylated forms of BMP6 have been reported from previous studies assaying BMP6 in serum and human brain hippocampus (31, 32). Since BMP6 can bind the type II receptor irrespective of its glycosylation state (15, 24) and since ALK2 exists in preformed heterodimers with its partner type II receptors (33), it stands that unglycosylated BMP6 can still drive the formation of ALK2-ActRIIB heterotetramers. We predict that such heterotetramers would fail to signal and may act to reduce the overall level of signaling mediated by ALK2 without affecting signaling via ALK3. Furthermore, we speculate that this mechanism may be shared with other ALK2 ligands such as BMP5, BMP6, BMP7, and BMP8 which feature a conserved prehelical sequence and have been demonstrated to be glycosylated similarly to BMP6 (SFig. 8).

Next, we explored the other class of ligands that interact with ALK2, the Activins, represented by Activin A. Given practical difficulties in structurally resolving an ALK2-ActA complex, we modeled ALK2 interaction with ActA based on the ALK2-BMP6 complex, using AlphaFold. The model revealed a key interaction between the ALK2 β4β5 loop and the fingertip of ActA, which we verified experimentally. Furthermore, we demonstrate that upon ligand binding, ALK2 undergoes a conformational change in the β4β5 loop that is greater than that seen with other BMP type I receptors, positioning ALK2 closer to the ligand fingers. This difference is likely due to β4β5 loop having additional amino acids in the N-terminal half of the β4β5 loop when compared to ALK1, ALK3, or ALK6 (Fig. 2D). A similar extension is seen in the activin receptors, ALK4 and ALK5 but not ALK7. Our data show that while this altered position of the β4β5 loop is not critical for BMP6, it plays roles in ActA binding. Here, modulation of this prospective interface by mutagenesis of residues from either the receptor (ALK2.Y74E) or the ligand (ActA.deltaD406) completely ablates the interaction. Interestingly, this fingertip interaction is similar to what is observed during ALK4:ActA interaction, where the ligand fingertip residue D406 directly binds to ALK4.

Sequence alignment of ALK2 across a variety of species reveals a widespread conservation of the critical residue for ActA interaction, Tyr^74^. Interestingly, in zebrafish and drosophila there is a proline in this position and in salmon, a serine (SFig. 8). The ALK2 homolog in drosophila, Saxophone (Sax), has been demonstrated to form nonsignaling complexes that sequester BMP homologs (46). Furthermore, these complexes are similarly converted to signaling complexes when a FOP-homologous mutation, K262H, is introduced to Sax (47). However, overexpression of Sax has no effect on either ActA or ActB homologs (Dawdle and dActB), confirming a lack of interaction. This is consistent with experiments in zebrafish that display increased BMP signaling with FOP mutants, but not due to ActA (34). Thus, it appears that while the ability to engage BMP ligands and form nonsignaling complexes is conserved between Sax and ALK2, interactions with the activin class are more evolutionarily recent, perhaps as a part of the evolution of load-bearing bones (4, 9, 35).

ALK2 remains one of the more perplexing TGFβ receptors, with interactions between two major branches of ligands, the BMPs and the Activins. The structural resolution of ALK2-ActRIIB:BMP6 and ALK3-ActRIIB:BMP6, and AlphaFold modeling of ALK2-ActRIIB:ActA reveals ALK2 as a ‘hybrid’ receptor, able to act similarly to a BMP type I receptor at the wrist interface or an activin type I receptor (such as ALK4) at the fingertip. Additionally, these structures have allowed us to directly compare how BMP6 binds to two distinct type I receptors highlighting the importance of the BMP6 post-translational modification for binding to ALK2. Future work will need to explore the role of the ALK2 nonsignaling complex as well as comprehensively characterize the various glycostates of BMP6 and their potential roles *in vivo*. Overall, our structures serve to not only to detail the interactions between ALK2 and its ligands, but to also advance our molecular understanding of TGFβ type I receptors and the mechanisms of specificity used to dictate each unique receptor:ligand combination.

## Methods

### Hetero– and Homodimeric Receptor Trap Expression and Purification

ActRIIB_2-_Fc along with ALK2-Fc, ALK3-Fc and the corresponding mutant ALK2/3-Fc fusions were expressed and purified from Chinese hamster ovary (CHO) cells as previously described (17, 19). The heterodimeric traps were also produced and purified from CHO cells. Here, constructs containing each receptor-Fc fusion were designed to favor heterodimer formation through either the Fc-Star strategy (ALK2-ActRIIB-Fc) or a knob-in-hole strategy (ALK3-ActRIIB-Fc) (36, 37). To express each molecule, plasmids were co-transfected at a ratio of 60% ActRIIB to 40% type I receptor fusion into CHO cells. Following 10 days of expression, conditioned media was harvested, centrifuged and filtered prior to loading through a MabSelect SuRe protein A column (Cytiva). Here, ALK2-ActRIIB-Fc was eluted through a pH gradient from pH 7.5 (20mM Tris, 300mM NaCl) to pH 3 (20mM Sodium Citrate, 300mM NaCl) to separate contaminating homodimeric ALK2_2_-Fc. ALK3-ActRIIB-Fc was eluted in a batch manner. Semi-purified protein was then further purified by size exclusion chromatography (SEC) using a Superdex 200 increase column (Cytiva).

For structural studies, ALK2-ActRIIB-Fc and ALK3-ActRIIB-Fc were then digested with IdeS-His protease at 37°C overnight, cleaving the Fc portion of each construct below the antibody hinge. The cleaved Fc and IdeS-His were then removed from ALK2-ActRIIB and ALK3-ActRIIB by both CaptureSelect (Thermo) and TALON (Cytiva) resins, respectively. To further purify the digested receptor fusions, the sample was loaded on a Superdex 200 increase column (Cytiva). 20mM Tris 7.5, 300mM NaCl was used as the running buffer for both resins and the final polishing SEC.

Human recombinant ActA was produced to homogeneity as previously described (5). Recombinant human BMP6 was purchased from RnD (Cat. No. 507-BP), where it was produced from CHO cells. For applicable data, BMP6 was deglycosylated with PNGase F (New England Biolabs) overnight and digestion was confirmed through SDS-page analysis.

### CryoEM Sample Preparation and Data Collection

Digested and purified ALK2-ActRIIB and ALK3-ActRIIB were mixed with BMP6 at a 2.1:1 receptor trap:ligand molar ratio and incubated at room temperature for 30 min. Complexes were purified through SEC on a Superdex 200 increase column (Cytiva). For each complex, distinct peaks consistent with expected complex molecular weights were observed and pooled for subsequent concentration for structural studies.

CryoEM grids of both purified complexes of ALK2-ActRIIB:BMP6 and ALK3-ActRIIB:BMP6 were prepared at a protein concentration of ∼3mg/mL. Samples were supplemented with PMAL-C8 amphipol (Anatrace) to a final concentration of 0.15% immediately prior to grid preparation to aid in vitrification. UltrAuFoil 1.2/1.3 grids were used for both protein samples following plasma cleaning in a Solarus II (Gatan) using a H_2_O_2_ gas mixture. A Vitrobot Mark IV (Thermo Fisher) operated at 4°C and 70% humidity was used for blotting the grids and plunge freezing them into liquid ethane cooled by liquid nitrogen.

Grids were loaded into a Titan Krios G3i electron microscope equipped with a Bioquantum K3 (Gatan). Images were collected in counted mode at a nominal magnification of 105,000x, yielding a pixel size of 0.839 Å. For ALK2-ActRIIB:BMP6 and ALK3-ActRIIB:BMP6, defocus ranges of – 0.8 to –2.4 µm and –1.0 to –3.4 µm, respectively, were set for data collection in EPU (ThermoFisher). The energy filter was inserted with slit width 20 eV. Each movie was dose fractionated into 46 frames over a 2 second exposure and had a total dose of ∼40 electrons per Å^2^. Further details of the data collections leading to the structures are shown in Supplemental Table 2.

### CryoEM Data Processing

CryoEM data were processed using cryoSPARC v4, where the same general workflow was used for each sample(38). Exact workflows and details for each structure are available in Supplementary Figure 3 and the statistics, in Supplemental Table 2. In summary, movies were motion and CTF corrected using cryoSPARC’s patch implementations. Micrographs with poor resolution estimates were removed from further processing. Particles were then blob-picked followed by multiple rounds of 2D classification to generate templates for particle picking and subsequent 2D and 3D classification. Once a subset of particles was obtained that resulted in a homogenous population and 3D volume, a topaz model was trained against particle coordinates from a subset of the initial exposures(39). The model was then used to pick particles across the entire dataset, which were subject to multiple rounds of 2D and 3D classification. Finally, maps were refined sequentially against alternative classes (Heterogeneous refinement) to further increase particle homogeneity and ultimately, C2 symmetry was applied to refinement, in the case of ALK2-ActRIIB:BMP6 and ALK3-ActRIIB:BMP6(40). Sharpened and unsharpened maps were carried forward for visualization and model building.

### Model Building and Refinement

Manual model building was conducted in Coot 0.9.6 and real space refinement of models was conducted using Phenix 1.21(41, 42). The initial model for building ALK2-ActRIIB’:BMP6 was constructed by combining ALK2 (PDB code: 7YRU), ActRIIB (PDB: 7MRZ) and BMP6 (2QCW). Each individual component was aligned to ALK3:ActRIIA:BMP2. This structure was then used as a starting model for building ALK3-ActRIIB’BMP6 after ALK3 (PDB code: 2GOO) was aligned in place of ALK2(23). PyMOL and USCF Chimera X were used to visualize models and maps(43). Buried surface area and subsequent related calculations were carried out in Phenix 1.21-Pisa (42). The model of ALK2 bound to ActA was generated utilizing AlphaFold 2 (32). Five AMBER relaxed complex models were generated from input sequences featuring the ALK2-ECD and ActA-mature domain and the highest ranked model was used for analysis.

### Surface Plasmon Resonance

SPR affinity analysis was performed on a Cytiva T-200 instrument using T200 Control Software version 3.2.1 (Cytiva) for data acquisition. A CM4 sensor chip (Cytiva) was prepared by EDC/NHS immobilization of a goat antihuman Fc-specific IgG (in-house) of approximately 3000RU. Experiments were carried out in HBS-EP+ buffer (10mM Hepes pH 7.4, 350mM NaCl, 3.4mM EDTA, 0.05% P-20 surfactant) at 25°C. A 5-step, twofold serial dilution was performed in the aforementioned buffer for each ligand, with an initial concentration of 5nM for the studies with ActRIIB_2_-Fc, ALK2-ActRIIB-Fc, ALK3-ActRIIB-Fc, and ActRIIB_1_-Fc and 20nM for the studies with ALK2-Fc, ALK3-Fc and the corresponding mutants. Each cycle had a ligand association and dissociation time of 300s. The flow rate for kinetics was maintained at 30uL/min. SPR chips were regenerated with 20mM H_3_PO_4_ pH 2.5. Kinetic analysis was conducted using the Biacore T200 evaluation software using a 1:1 fit model with mass transport limitation (red lines).

### Luciferase Reporter Assays

Assays using the BRE (BMP responsive element) luciferase reporter cells were performed in a manner similar to those previously described(4). Briefly, cells were plated in a 96-well format (3 x 10^4^ cells/well) and propagated for 24hr. For EC50 experiments (Fig. 4C), growth media was replaced with serum free media supplemented with 0.1% BSA and the desired ligand, where a threefold serial dilution was performed with a starting concentration of 54nM (ActA and BMP6). Cells were incubated for 18 hr before lysing and assaying for luminescence using a Victor X Light Multilabel Reader (Perkin Elmer, Waltham, MA). For the assays featuring stable expression of ALK2 and corresponding mutants, cell-lines were generated through lentiviral transduction similar to previous work using reagents obtained from Origene(44). Here, constructs containing the full-length ALK2 receptor were designed in a pLVX vector backbone and co-transfected with lentiviral packaging plasmids into HEK293T cells. Viral particles were harvested 48hr later and used to transduce BRE luciferase reporter cells. Following selection of stable pools with hygromycin, cells were sorted for expression utilizing antibodies against ALK2 (Mouse; Mab637; RnD) and Mouse IgG (Donkey Alexa Fluor 647 conjugated; A-31571; Life Technologies). The activity data were imported into GraphPad Prism and fit using a non-linear regression to calculate the EC_50_ or IC_50_.

## Figures, Tables and Supplemental Information (In order of appearance in manuscript)

**Supplemental Figure 1.**
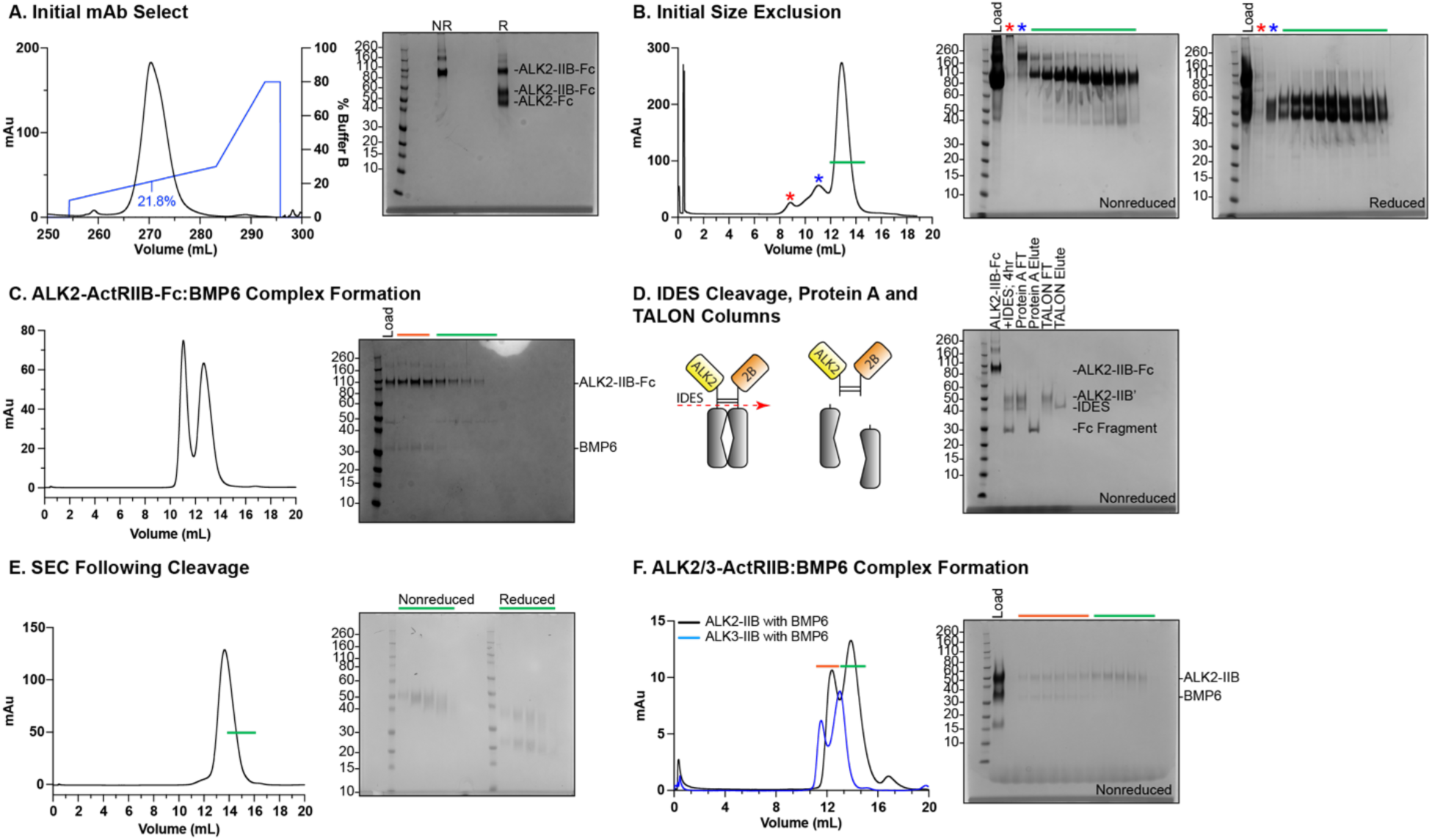
Representative biochemistry, purification, and complex formation of ALK2-ActRIIB:BMP6. **A.** Anti-Fc column (mAb Select) run to purify ALK2-ActRIIB-Fc from conditioned media. SDS PAGE gel displays non-reduced (NR) and reduced (R) samples from major peak. **B.** Size exclusion to purify ALK2-ActRIIB-Fc and corresponding nonreduced and reduced SDS PAGE gels. *Red* and *blue* Asterisks correspond to samples from respective peaks along with samples from throughout the major peak represented by the *green* bar. C. Complex formation and subsequent size exclusion of ALK2-ActRIIB-Fc:BMP6. Complex formation confirmed through SDS PAGE. D. IDES cleavage schematic and SDS PAGE gel displaying flowthrough (FT) and elution products following sequential Protein A and TALON gravity columns. **E.** Size exclusion to purify ALK2-ActRIIB to monodispersity. **F.** Complex formation and subsequent size exclusion between ALK2-ActRIIB (*black*). and ALK3-ActRIIB (*blue*) with BMP6. Samples on SDS PAGE gel correspond to ALK2-ActRIIB trace (*black*) and are from the corresponding peak with colored band above.

**Supplemental Figure 2.**
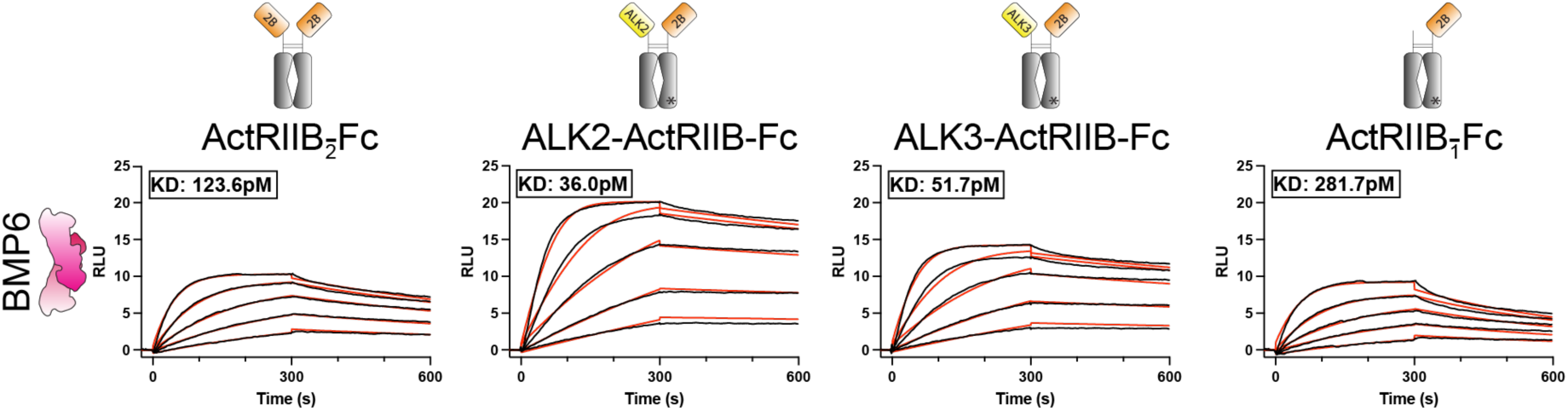
Heterodimeric traps, ALK2-ActRIIB-Fc and ALK3-ActRIIB-Fc bind BMP6 with high affinity. SPR sensorgrams of BMP6 binding to anti-human antibody-captured ActRIIB_2_-Fc, ALK2-ActRIIB-Fc, ALK3-ActRIIB-Fc, or ActRIIB_1_-Fc. Sensorgrams (*black lines*) are overlaid with fits to a 1:1 interaction model with mass transport limitations (*red lines*). Each experiment was performed in triplicate and the kinetic parameters are summarized in STable 1.

**Supplemental Table 1.** SPR Kinetic Analysis.

| Analyte | Ligand | $k_a$ ($M^{-1}s^{-1}$) | $k_d$ ($s^{-1}$) | $K_D$ (pM)<sup>a</sup> |
| --- | --- | --- | --- | --- |
| BMP6 | ActRIIB<sub>2</sub>-Fc | $1.03 \pm 0.007 \times 10^7$ | $1.27 \pm 0.03 \times 10^{-3}$ | $123 \pm 3.00$ |
| BMP6 | ALK2-ActRIIB-Fc | $1.38 \pm .02 \times 10^7$ | $4.96 \pm 0.27 \times 10^{-4}$ | $36.0 \pm 2.5$ |
| BMP6 | ActRIIB<sub>1</sub>-Fc | $8.23 \pm 1.26 \times 10^6$ | $2.29 \pm .008 \times 10^{-3}$ | $281.7 \pm 32.1$ |
| BMP6 | ALK3-ActRIIB-Fc | $1.27 \pm 0.02 \times 10^7$ | $6.46 \pm 0.65 \times 10^{-4}$ | $51.7 \pm 5.2$ |

<sup>a</sup>All kinetic parameters were analyzed using the Biacore T200 evaluation software using a 1:1 binding model.

Supplemental Table 1. SPR Kinetic Analysis.
| Analyte | Ligand | $k_a$ ($M^{-1}s^{-1}$) | $k_d$ ($s^{-1}$) | $K_D$ (pM)<sup>a</sup> |
| --- | --- | --- | --- | --- |
| dgBMP6 | ActRIIB<sub>2</sub>-Fc | $3.80 \pm 2.68 \times 10^6$ | $4.26 \pm 2.37 \times 10^{-4}$ | $132.7 \pm 44.0$ |
| dgBMP6 | ALK2-ActRIIB-Fc | $4.55 \pm 1.32 \times 10^6$ | $6.90 \pm 1.02 \times 10^{-4}$ | $164.3 \pm 65.5$ |
| dgBMP6 | ActRIIB<sub>1</sub>-Fc | $8.08 \pm 3.64 \times 10^6$ | $2.38 \pm 1.09 \times 10^{-3}$ | $293.7 \pm 3.51$ |
| dgBMP6 | ALK3-ActRIIB-Fc | $3.96 \pm 2.38 \times 10^6$ | $2.43 \pm 0.47 \times 10^{-4}$ | $71.4 \pm 25.6$ |

<sup>a</sup>All kinetic parameters were analyzed using the Biacore T200 evaluation software using a 1:1 binding model.

**Supplemental Figure 3.**
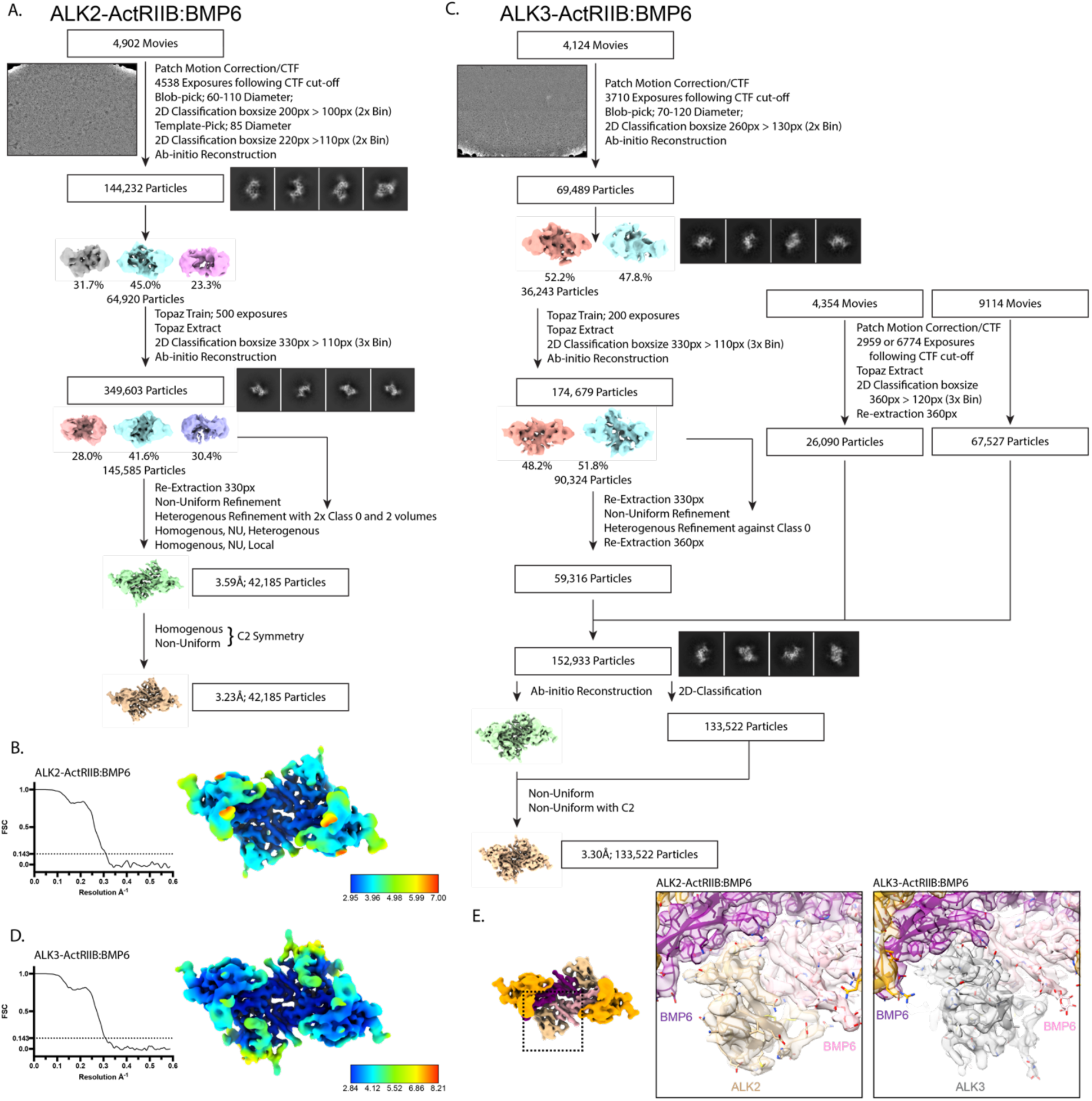
CryoEM data processing and reconstruction of BMP6 in complex with ALK2-ActRIIB and ALK3-ActRIIB. **A.** Data processing flow chart for ALK2-ActRIIB:BMP6. **B.** FSC curve output and local resolution map by cryoSPARC for ALK2-ActRIIB:BMP6. **C.** Data processing flow chart for ALK3-ActRIIB:BMP6. **D.** FSC curve output and local resolution map by cryoSPARC for ALK3-ActRIIB:BMP6. **E.** Overlay of sharpened maps and model focused on the respective type I receptors.

**Supplemental Table 2.** CryoEM data, structure refinement and validation.

|  | ALK2-ActRIIB:BMP6 (XXX) | ALK3-ActRIIB:BMP6 (XXX) |
| --- | --- | --- |
| **Data Collection and Processing** |  |  |
| Magnification | 105,000 | 105,000 |
| Voltage (kV) | 300 | 300 |
| Electron exposure (e<sup>-</sup>/Å<sup>2</sup>) | ~40 | ~40 |
| Defocus Range (μm) | -0.8 to -2.4 | -1 to -3.4 |
| Pixel size (Å) | 0.839 | 0.839 |
| Number of Movies | 4902 | 13,443 (3710 + 2959 + 6774) |
| Initial number of particles | 1,820,762 | 3,218,335 (778,719 + 752,555 + 1,686,061) |
| Particles selected after 2D classification | 349,603 | 152,933 (59,316 + 26,090 + 67,527) |
| Final selected particles | 42,185 | 133,522 |
| Symmetry imposed | C2 | C2 |
| Map resolution (Å) | 3.2 | 3.3 |
| FSC threshold | 0.143 | 0.143 |
| **Refinement** |  |  |
| Initial Model used | PDBs: 6MAC, 6OMO, 7YRU | PDBs: 2GOO, 6MAC, 6OMO |
| **Model Composition** |  |  |
| Non-hydrogen atoms | 4578 | 4822 |
| Protein Residues | 552 | 576 |
| Ligands | BMA: 4; NAG: 10 | BMA: 4, NAG: 14 |
| **R.m.s Deviations** |  |  |
| Bond lengths (Å) | 0.003 | 0.003 |
| Bond Angles (°) | 0.759 | 0.726 |
| **Validation** |  |  |
| MolProbity Score | 1.67 | 1.73 |
| Rotamer Outliers (%) | 0.00 | 0.39 |
| Clash Score | 4.44 | 5.39 |
| **Ramachandran Plot** |  |  |
| Favored (%) | 92.96 | 93.26 |
| Allowed (%) | 7.04 | 6.74 |
| Disallowed (%) | 0.00 | 0.00 |
| **Deposition ID** |  |  |
| PDB | XXX | XXX |
| EMDB | XXX | XXX |

**Supplemental Figure 4.**
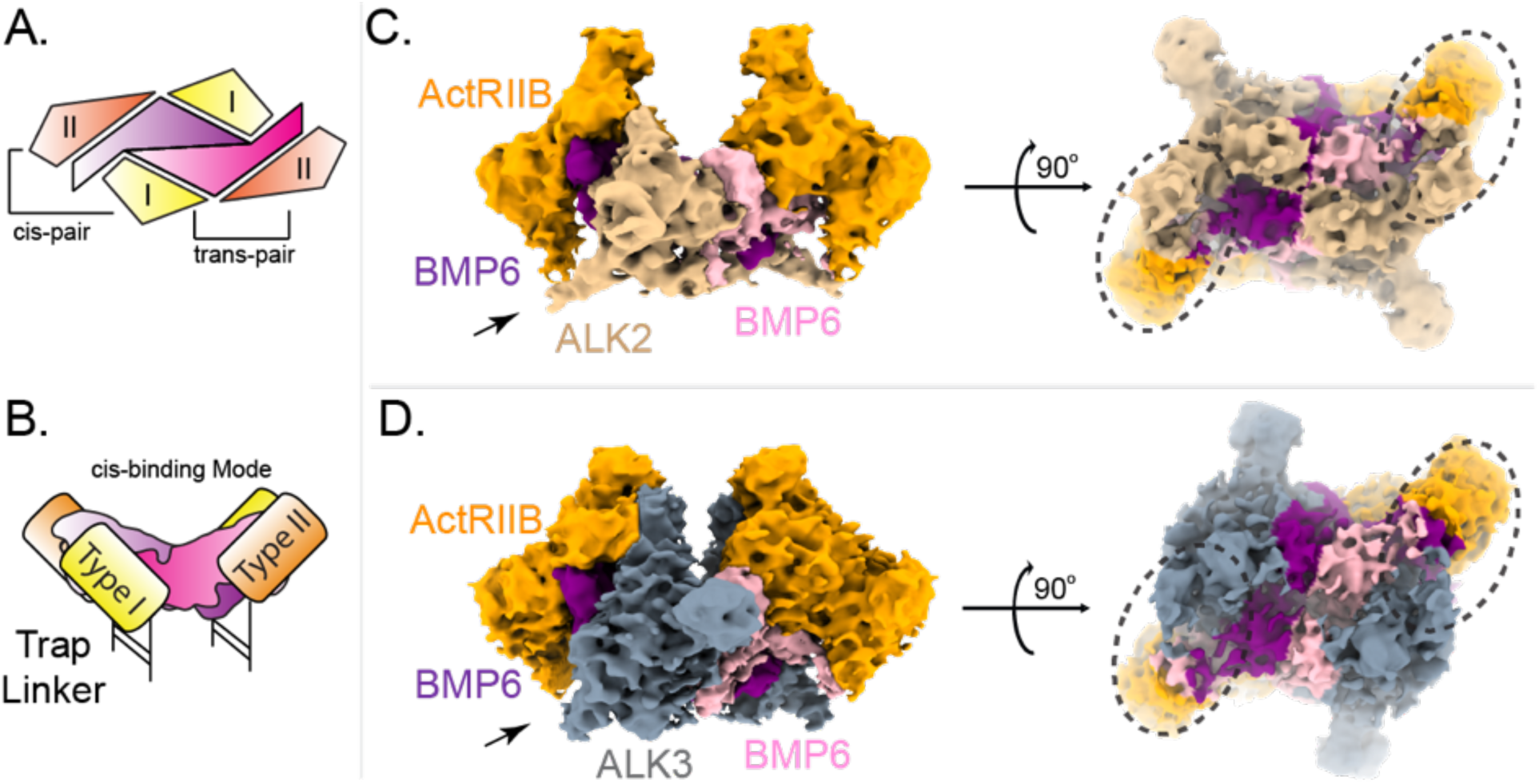
ALK2-ActRIIB and ALK3-ActRIIB bind BMP6 in a cis-manner. **A.** Schematic displaying two the two potential pairs of receptors in the heterotetrametric complex. **B.** Schematic displaying the Fc-cleaved Trap binding ligand in a cis-binding mode. **C. and D.** CryoEM maps displaying weakly resolved signal (indicated by black arrows) for intact antibody-hinge tether between the cis-pair of receptors in ALK2-ActRIIB:BMP6 (**C.**) and ALK3-ActRIIB:BMP (**D.**).

**Supplemental Figure 5.**
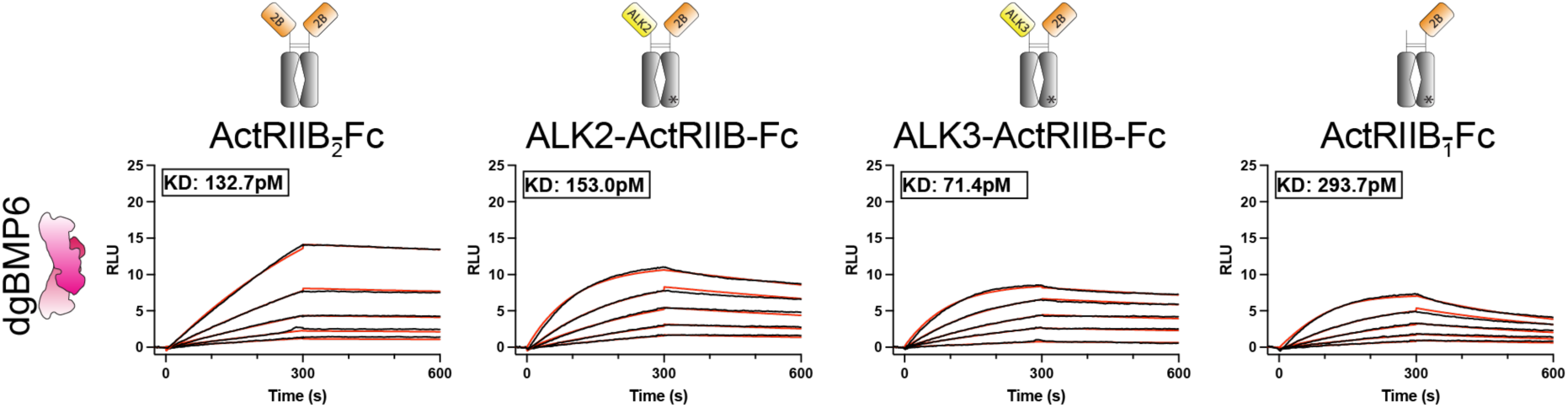
Deglycosylation of BMP6 decreases affinity for ALK2-ActRIIB-Fc. SPR sensorgrams of deglycosylated BMP6 (dgBMP6) binding to anti-human antibody-captured ActRIIB_2_-Fc, ALK2-ActRIIB-Fc, ALK3-ActRIIB-Fc, or ActRIIB_1_-Fc. Sensorgrams (*black lines*) are overlaid with fits to a 1:1 interaction model with mass transport limitations (*red lines*). Each experiment was performed in triplicate and the kinetic parameters are summarized in Supplemental Table 1.

**Supplemental Figure 6.**
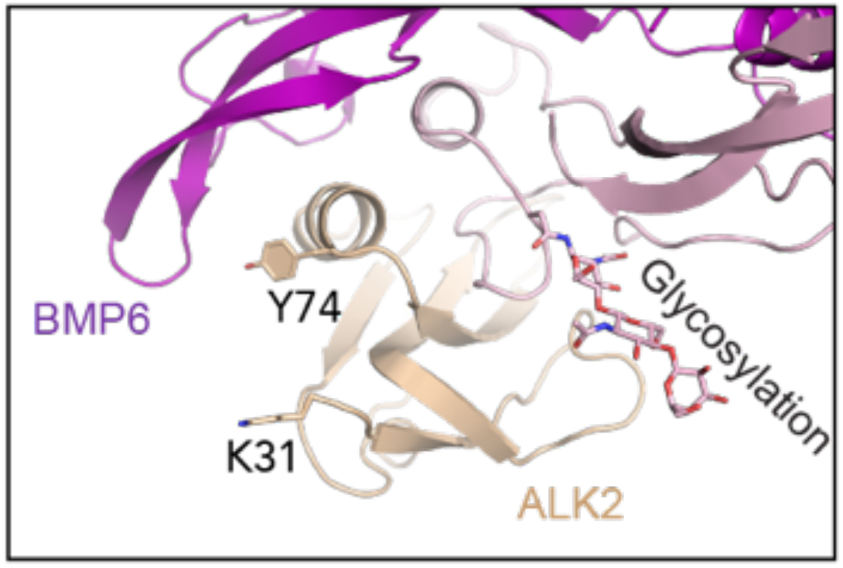
Structural zoom into ALK2-ActRIIB:BMP6 that displays the positioning of residues Tyr^74^ and Lys^31^, previously modeled to interact with the glycosylation on BMP6 (15).

**Supplemental Figure 7.**
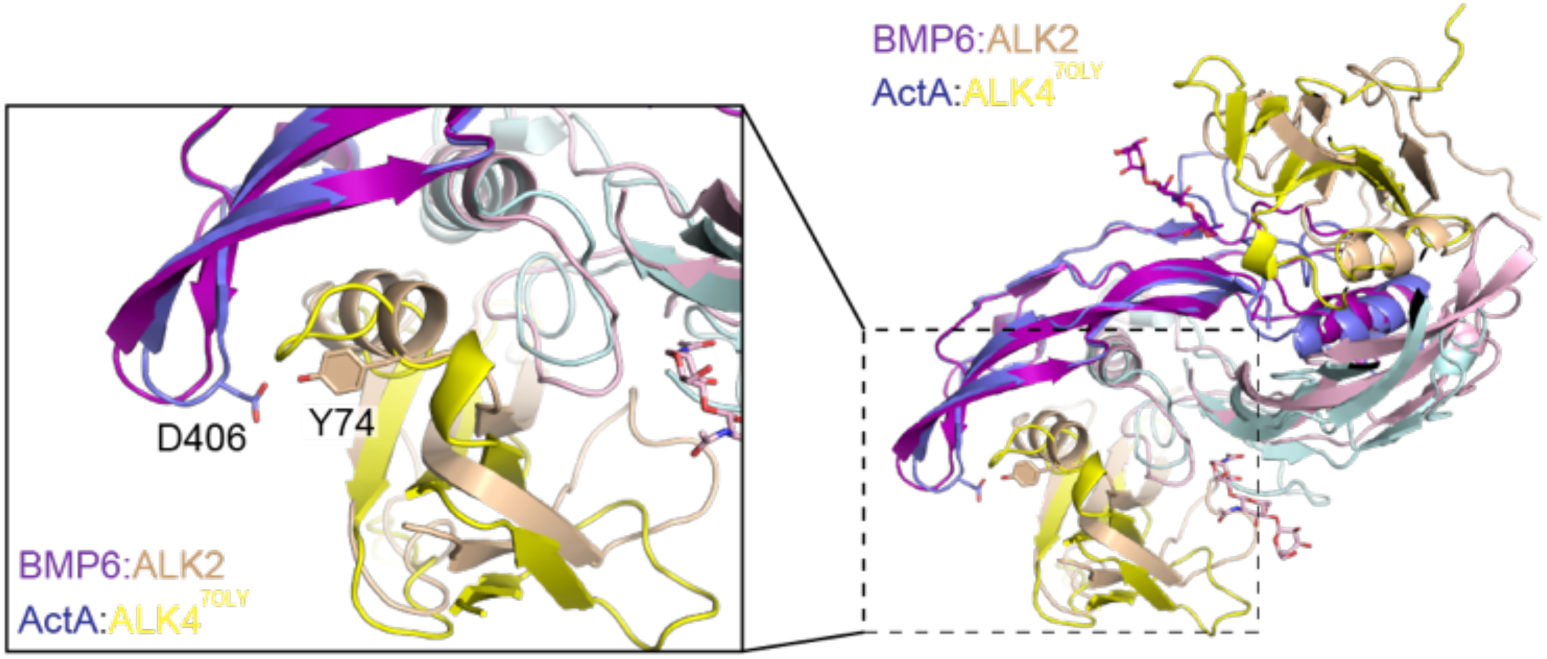
Alignment of structures: ALK2-ActRIIB:BMP6 and ALK4-ActRIIB:ActA (PDB code: 7OLY (19)). ActRIIB is hidden in this representation and alignment performed on darker colored ligand monomers. Zoom in highlights positioning of the ALK2 β4β5 loop and residue, Y74.

**Supplemental Figure 8.**
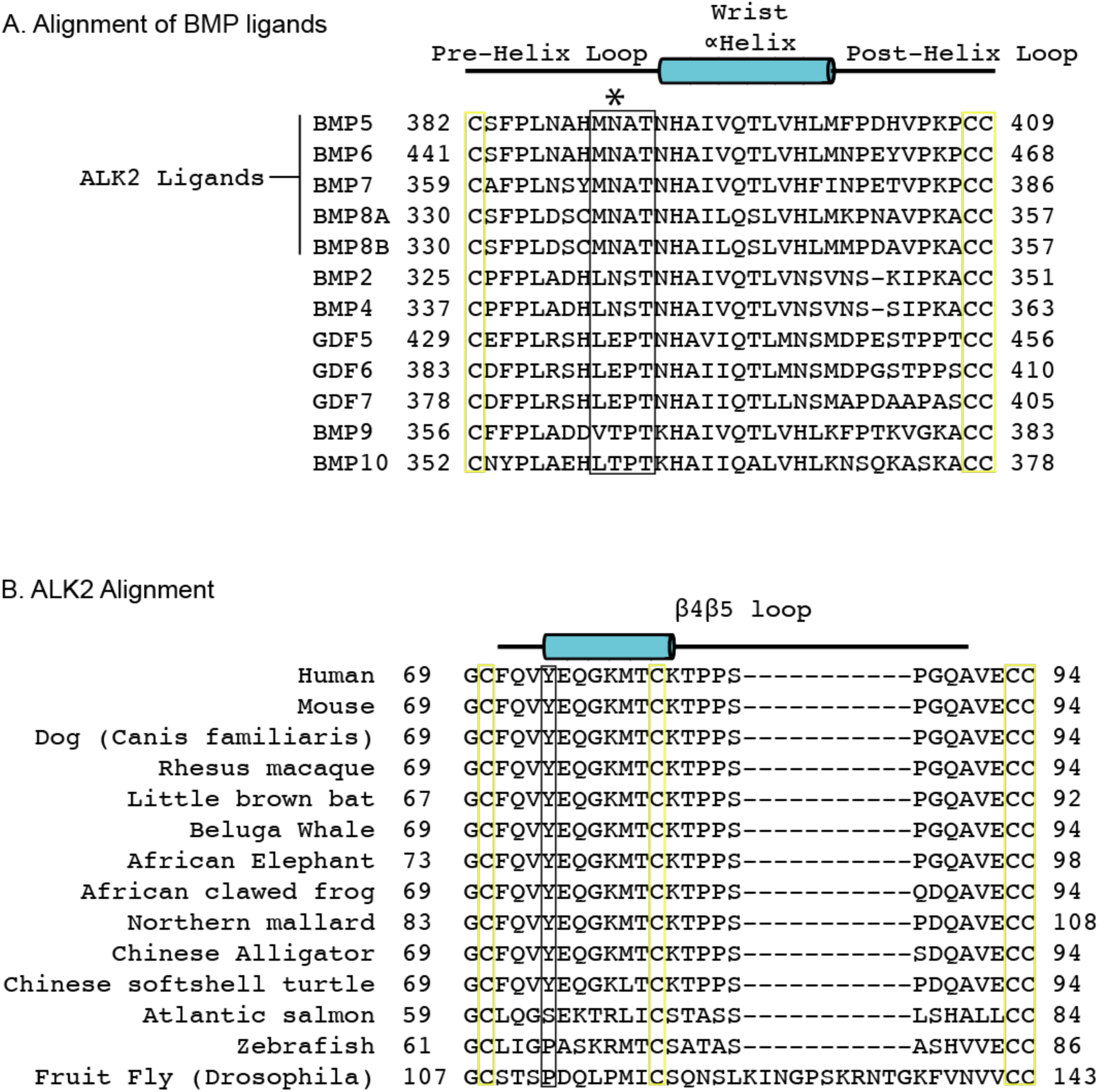
Sequence alignment of BMP wrist-helix regions and ALK2 across Species. A. Alignment of human BMP ligands, highlighting the wrist helix region. Asterisk denotes N-linked glycosylation site, while flanking residues are boxed in *black.* Conserved cystine positions flanking the wrist helix region are boxed in *yellow*. B. Alignment of the β4β5 loop region from the sequences of several species. ActA-critical and well-conserved residue, Tyr^74^ boxed in *black*.

## Acknowledgements.

The authors would like to thank Drs. Ashique Rafique and Drew Dudgeon, as well as Lisa Chen for their assistance in Surface Plasmon Resonance Data Collection.

## Author contributions

E.J.G. designed and performed research, analyzed and prepared data and wrote the manuscript. S.A., K.S., V.J.I, M.C.F., and A.N.E. provided analysis, mentorship, training, research environment, and edited the manuscript.

## Data Availability

Data Deposition: The structures of ALK2-ActRIIB:BMP6 and ALK3-ActRIIB:BMP6 described in this paper have been deposited to both the Protein Data Bank (XXX and XXX, respectively) and Electron Microscopy Data Bank (XXX and XXX, respectively).

## References.

1. A. P. Hinck, T. D. Mueller, T. A. Springer, Structural Biology and Evolution of the TGF-β Family. Cold Spring Harb. Perspect. Biol. 8, a022103 (2016).

2. E. J. Goebel, et al., Structural characterization of an activin class ternary receptor complex reveals a third paradigm for receptor specificity. Proc National Acad Sci 116, 15505–15513 (2019).

3. Y. E. Antebi, et al., Combinatorial Signal Perception in the BMP Pathway. Cell 170, 1184–1196.e24 (2017).

4. S. J. Hatsell, et al., ACVR1R206H receptor mutation causes fibrodysplasia ossificans progressiva by imparting responsiveness to activin A. Sci. Transl. Med. 7, 303ra137 (2015).

5. S. Aykul, et al., Activin A forms a non-signaling complex with ACVR1 and type II Activin/BMP receptors via its finger 2 tip loop. Elife 9, e54582 (2020).

6. K. Tsuchida, L. S. Mathews, W. W. Vale, Cloning and characterization of a transmembrane serine kinase that acts as an activin type I receptor. Proc. Natl. Acad. Sci. 90, 11242–11246 (1993).

7. L. Attisano, et al., Identification of human activin and TGFβ type I receptors that form heteromeric kinase complexes with type II receptors. Cell 75, 671–680 (1993).

8. M. Macías-Silva, P. A. Hoodless, S. J. Tang, M. Buchwald, J. L. Wrana, Specific Activation of Smad1 Signaling Pathways by the BMP7 Type I Receptor, ALK2*. J. Biol. Chem. 273, 25628–25636 (1998).

9. K. Hino, et al., Neofunction of ACVR1 in fibrodysplasia ossificans progressiva. Proc. Natl. Acad. Sci. 112, 15438–15443 (2015).

10. O. E. Olsen, et al., Activin A inhibits BMP-signaling by binding ACVR2A and ACVR2B. Cell Commun. Signal. 13, 27 (2015).

11. W. W. Hom, et al., Lysosomal degradation of ACVR1-Activin complexes negatively regulates signaling of Activins and Bone Morphogenetic Proteins. bioRxiv 2024.01.29.577837 (2024). 10.1101/2024.01.29.577837.

12. S. Aykul, E. Martinez-Hackert, Determination of half-maximal inhibitory concentration using biosensor-based protein interaction analysis. Anal. Biochem. 508, 97–103 (2016).

13. K. Nolan, et al., Structure of Gremlin-2 in Complex with GDF5 Gives Insight into DAN-Family-Mediated BMP Antagonism. Cell Rep. 16, 2077–2086 (2016).

14. S. A. Townson, et al., Specificity and Structure of a High Affinity Activin Receptor-like Kinase 1 (ALK1) Signaling Complex. J. Biol. Chem. 287, 27313–27325 (2012).

15. S. Saremba, et al., Type I receptor binding of bone morphogenetic protein 6 is dependent on N-glycosylation of the ligand. FEBS J. 275, 172–183 (2008).

16. A. N. Economides, et al., Cytokine traps: multi-component, high-affinity blockers of cytokine action. Nat. Med. 9, 47–52 (2003).

17. M. Humbert, et al., Sotatercept for the Treatment of Pulmonary Arterial Hypertension. N. Engl. J. Med. 384, 1204–1215 (2021).

18. J. Li, et al., ActRIIB:ALK4-Fc alleviates muscle dysfunction and comorbidities in murine models of neuromuscular disorders. J. Clin. Investig. 131, e138634 (2021).

19. E. J. Goebel, et al., Structures of activin ligand traps using natural sets of type I and type II TGFβ receptors. iScience 25, 103590 (2022).

20. E. J. Goebel, K. N. Hart, J. C. McCoy, T. B. Thompson, Structural biology of the TGFβ family. Exp. Biol. Med. 244, 1530–1546 (2019).

21. G. R. Gipson, et al., Structural perspective of BMP ligands and signaling. Bone 140, 115549 (2020).

22. E. J. Goebel, et al., Structural characterization of an activin class ternary receptor complex reveals a third paradigm for receptor specificity. Proc National Acad Sci 116, 15505–15513 (2019).

23. G. P. Allendorph, W. W. Vale, S. Choe, Structure of the ternary signaling complex of a TGF-β superfamily member. Proc. Natl. Acad. Sci. 103, 7643–7648 (2006).

24. H. J. Seeherman, et al., A BMP/activin A chimera is superior to native BMPs and induces bone repair in nonhuman primates when delivered in a composite matrix. Sci. Transl. Med. 11 (2019).

25. S. Arndt, et al., Iron-Induced Expression of Bone Morphogenic Protein 6 in Intestinal Cells Is the Main Regulator of Hepatic Hepcidin Expression In Vivo. Gastroenterology 138, 372–382 (2010).

26. L. Crews, et al., Increased BMP6 Levels in the Brains of Alzheimer’s Disease Patients and APP Transgenic Mice Are Accompanied by Impaired Neurogenesis. J. Neurosci. 30, 12252– 12262 (2010).

27. A. Kotzsch, J. Nickel, A. Seher, W. Sebald, T. D. Müller, Crystal structure analysis reveals a spring-loaded latch as molecular mechanism for GDF-5–type I receptor specificity. EMBO J. 28, 937–947 (2009).

28. D. Weber, et al., A silent H-bond can be mutationally activated for high-affinity interaction of BMP-2 and activin type IIB receptor. BMC Struct. Biol. 7, 6 (2007).

29. A. Kotzsch, et al., Structure Analysis of Bone Morphogenetic Protein-2 Type I Receptor Complexes Reveals a Mechanism of Receptor Inactivation in Juvenile Polyposis Syndrome*. J. Biol. Chem. 283, 5876–5887 (2008).

30. J. Guo, et al., Crystal structures of BMPRII extracellular domain in binary and ternary receptor complexes with BMP10. Nat. Commun. 13, 2395 (2022).

31. R. M. Salmon, et al., Molecular basis of ALK1-mediated signalling by BMP9/BMP10 and their prodomain-bound forms. Nat. Commun. 11, 1621 (2020).

32. J. Jumper, et al., Highly accurate protein structure prediction with AlphaFold. Nature 596, 583–589 (2021).

33. S. Aykul, et al., Anti-ACVR1 antibodies exacerbate heterotopic ossification in fibrodysplasia ossificans progressiva (FOP) by activating FOP-mutant ACVR1. J. Clin. Investig. 132, e153792 (2022).

34. B. E. Mucha, M. Hashiguchi, J. Zinski, E. M. Shore, M. C. Mullins, Variant BMP receptor mutations causing fibrodysplasia ossificans progressiva (FOP) in humans show BMP ligand-independent receptor activation in zebrafish. Bone 109, 225–231 (2018).

35. E. M. Shore, et al., A recurrent mutation in the BMP type I receptor ACVR1 causes inherited and sporadic fibrodysplasia ossificans progressiva. Nat. Genet. 38, 525–527 (2006).

36. J. B. B. Ridgway, L. G. Presta, P. Carter, ‘Knobs-into-holes’ engineering of antibody CH3 domains for heavy chain heterodimerization. *Protein Eng.*, Des. Sel. 9, 617–621 (1996).

37. A. D. Tustian, C. Endicott, B. Adams, J. Mattila, H. Bak, Development of purification processes for fully human bispecific antibodies based upon modification of protein A binding avidity. mAbs 8, 828–838 (2016).

38. A. Punjani, J. L. Rubinstein, D. J. Fleet, M. A. Brubaker, cryoSPARC: algorithms for rapid unsupervised cryo-EM structure determination. Nat. Methods 14, 290–296 (2017).

39. T. Bepler, K. Kelley, A. J. Noble, B. Berger, Topaz-Denoise: general deep denoising models for cryoEM and cryoET. Nat. Commun. 11, 5208 (2020).

40. A. Punjani, H. Zhang, D. J. Fleet, Non-uniform refinement: adaptive regularization improves single-particle cryo-EM reconstruction. Nat. Methods 17, 1214–1221 (2020).

41. P. Emsley, B. Lohkamp, W. G. Scott, K. Cowtan, Features and development of Coot. Acta Crystallogr. Sect. D: Biol. Crystallogr. 66, 486–501 (2010).

42. D. Liebschner, et al., Macromolecular structure determination using X-rays, neutrons and electrons: recent developments in Phenix. Acta Crystallogr. Sect. D, Struct. Biol. 75, 861–877 (2019).

43. E. F. Pettersen, et al., UCSF ChimeraX: Structure visualization for researchers, educators, and developers. Protein Sci. 30, 70–82 (2020).

44. J. Elegheert, et al., Lentiviral transduction of mammalian cells for fast, scalable and high-level production of soluble and membrane proteins. Nat. Protoc. 13, 2991–3017 (2018).

